# Close encounters between infants and household members measured through wearable proximity sensors

**DOI:** 10.1101/232355

**Authors:** Laura Ozella, Francesco Gesualdo, Michele Tizzoni, Caterina Rizzo, Elisabetta Pandolfi, Ilaria Campagna, Alberto Eugenio Tozzi, Ciro Cattuto

## Abstract

Describing and understanding close proximity interactions between infant and family members can provide key information on transmission opportunities of respiratory infections within households. Among respiratory infections, pertussis represents a public health priority. Pertussis infection can be particularly harmful to young, unvaccinated infants and for these patients, family members represent the main sources of transmission. Here, we report on the use of wearable proximity sensors based on RFID technology to measure face-to-face proximity between family members within 16 households with infants younger than 6 months for 2-5 consecutive days of data collection. The sensors were deployed over the course of approximately 1 year, in the context of a national research project aimed at the improvement of infant pertussis prevention strategies. We recorded 5,958 contact events between 55 individuals: 16 infants, 4 siblings, 31 parents and 4 grandparents. The contact networks showed a heterogeneous distribution of the cumulative time spent in proximity with the infant by family members. Most of the contacts occurred between the infant and other family members (70%), and many contacts were observed between infants and adults, in particular between infant and mother, followed by father, siblings and grandparents. A larger number of contacts and longer contact durations between infant and other family members were observed in families adopting exclusive breastfeeding, compared to families in which the infant receives artificial or mixed feeding.

Our results demonstrate how a high-resolution measurement of contact matrices within infants’ households is feasible using wearable proximity sensing devices. Moreover, our findings suggest the mother is responsible for the large majority of the infant’s contact pattern, thus being the main potential source of infection for a transmissible disease. As the contribution to the infants’ contact pattern by other family members is very variable, vaccination against pertussis during pregnancy is probably the best strategy to protect young, unvaccinated infants.

## Introduction

Strategies to prevent infectious diseases transmitted through physical contact are informed by mathematical models describing disease transmission. Such models help identifying effective control measures and allow to predict their impact [1, 2]. Among the determinants included in disease transmission models, contact patterns (in particular age-specific contact patterns) are crucial to obtain a good estimation of transmission parameters [3]. Most of the studies on this issue have approximated contact matrices to data obtained through diary-based surveys. This approach permits obtaining only an approximate estimate of the number of contacts between individuals and of their duration, and may imply a substantial recall bias [4, 5].

Recent technological advances allow to measure real-world interactions using mobile devices and wearable sensors, in a variety of contexts, at very different spatial and temporal scales [6, 7]. Wearable sensors can gather data on human proximity and social interactions in an objective way and by means of non-obtrusive methodologies. With regard to infectious disease epidemiology, active Radio-Frequency IDentification (RFID) technology has been used to measure contact patterns in different settings relevant to the transmission of infectious agents: a paediatric hospital [8], a tertiary care hospital [9, 10], and a primary school [11].

Young infants represent a high risk group for transmissible infectious diseases. Among respiratory infections, pertussis can be particularly harmful for young, unvaccinated infants, that have a higher morbidity and mortality compared to other age groups [12-14]. Information on contact patterns within households can be useful to plan prevention strategies for protecting young infants from pertussis and other respiratory infections: infants <6 months mainly conduct an indoor life, and social interactions mainly happen within family members, who represent the main source of transmission [15-17], despite contacts with non-household members may play an important role [18]. Moreover, household interaction may vary among families, and specific caregiving practices, like breastfeeding, may be associated with a different contact pattern and, consequently, with a different risk of infectious disease transmission.

In this study, we report on the use of wearable proximity sensors based on RFID technology to measure face-to-face proximity and pattern of contacts between family members, within households with infants younger than 6 months. The main objectives of the study were: (1) to demonstrate the feasibility of contact measures through proximity sensing devices in the household setting; (2) to assess the differences in contact patterns between family members, as well as their structural and temporal heterogeneities; (3) to investigate whether socio-demographic variables or feeding practices affect contact patterns between parents and infants.

## Materials and Methods

### Study setting

The present study was conducted in the context of a national research project, funded by the Italian Ministry of Health, aimed at the improvement of infant pertussis prevention strategies. The study was approved by the Bambino Gesù Children’s Hospital Ethical Committee (protocol n. 866LB), and was conducted in Rome, Italy, from March 2015 to January 2016. Twenty households with a healthy infant younger than 6 months were selected from the list of patients of a family paediatrician. A household was defined as the group of people living in the same house of the infant. In a first, preliminary phase, members of each household were invited to a meeting in which the details and the aims of the study were explained. In this context, a signed informed consent was obtained from parents and relatives. All participants were given a Radio-Frequency IDentification (RFID) badge (see below) and were asked to wear it every time they were at home. The sensors were enclosed in a pouch with a lanyard and given to each participant, to be worn around the neck. Infants’ pouches were secured to a pacifier holder. Participants were asked to mark the times of entry and exit from the house during the entire study duration. Only data collected when the participants were inside the house were considered in the analysis. Each household received sensors on separate days and the family members wore the sensors for 2-5 days, depending on the household. On the last day of data collection, fieldworkers directly collected the sensors at the infants’ houses.

### Data collection

The data were obtained and processed using a proximity-sensing platform developed by the SocioPatterns collaboration [19]. The platform and its applications has been described in detail in several previous works [20-22]. The system is based on small active RFID devices (‘tags’) that exchange ultra-low-power radio packets in a peer-to-peer fashion, and perform a scan of their neighbourhood by alternating transmit and receive cycles. During the transmit phase, low-power packets are sent out on a specific radio channel; during the receive phase, the devices listen on the same channel for packets sent by nearby devices [8, 11, 20, 21]. When individuals wear the devices on their chest, exchange of radio packets between RFID devices is only possible when they are facing each other, as the human body acts as a RF shield at the carrier frequency used for communication. Sensors in close proximity exchange a maximum of about 1 power packet per second, and the exchange of low-power radio-packets is used as a proxy for the spatial proximity of the individuals wearing the sensors [8, 20]. In particular, the close proximity is measured by the attenuation, defined as the difference between the received and transmitted power. We defined that a ‘contact’ occurs between two individuals, during a time slice duration of 20 s, if, and only if, the RFID devices exchanged at least one radio packet during that interval. After a contact is established, it is considered ongoing as long as the devices continue to exchange at least one packet for every subsequent 20 s intervals. Conversely, a contact is considered broken if a 20 s interval elapses with no exchange of radio packets [11, 20]. Therefore, we set the attenuation threshold to detect proximity events between devices situated in the range 1 – 1.5 m of one another, thus corresponding to a close-contact situation, during which a communicable disease infection can be transmitted, either by droplet transmission through coughing or sneezing, or by direct physical contact [23]. Each device has a unique identification number that was used to link the information on the contacts established by the person carrying the device with his/her profile.

### Data analysis

#### Contact data and network analysis

We extracted and cleaned the data separately for each participant. Night contacts, collected between 9 pm and 7 am, were disregarded from the analysis, due to inconsistent spikes in data that suggested heightened interaction between family members, probably as a result of removal and storage of devices together. We computed the number of contact events recorded by each family member and the statistical distribution of the duration of contact events. The duration of contacts is expressed in seconds, minutes or hours depending on the context; the choice of the different time unit was made to increase readability and comprehension. We also generated aggregated contact networks of the full experimental time period. We considered individuals as nodes of the network, while the edges represented the presence of at least one recorded contact event between two individuals during the aggregation time window. Each network edge was weighted by the total time the two individuals spent in contact with each other during the aggregation time window. We measured the network density in each household, defined as the ratio of the number of observed edges to the number of potential edges.

Moreover, we analysed the temporal features of the aggregated contact networks. We compared the amount of time spent in proximity by two individuals, with respect to a circadian activity profile calculated for all the sensors of the experiment. We considered four different time fractions of the day: morning (from 7 am to 11 am), late morning - lunch (from 11 am to 2 pm), afternoon (from 2 pm to 5 pm), and late afternoon - evening (from 5 pm to 9 pm). Participants were grouped into four age categories [24]: <6 months old (infant), 1-5 (pre-school), 20-49 (adults), and > 50 (elderly) years old (enrolled families did not include infants aged 6 to 12 months).

Given a contact network, we defined as degree *k_i_* of a node *i* the number of distinct individuals with whom the individual *i* has been in contact, and the weight *w_ij_* of an edge between nodes *i* and *j* the cumulative duration of the contact events recorded between two individuals. Network edges are undirected and the weights on the edges are symmetric (*w_ij_* = *w_ji_*). We studied the statistical distributions of the degrees and weights of the contact networks. We generated contact matrices based on number and duration of contacts by age category. We also generated aggregated contact networks on a daily scale. In order to make the daily contact networks comparable across households, we considered only contacts collected over entire days (from 7 am to 9 pm).

To assess the changes in time duration of contacts of each household member, we compared the daily network structures by measuring the similarity between the neighbourhoods of each node across different days for each household. As each edge *i-j* in the aggregated network is weighted by the total time *i* and *j* spent in face-to-face proximity, the similarity between the neighbourhoods of an individual *i* in time *t_1_* and time *t_2_* can be quantified by the cosine similarity [11] defined as:

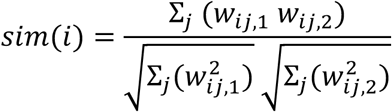

where *w_ij,1_* and *w_ij,2_* are the weights on the edge *i - j* measured at time *t_1_* and time *t_2_,* respectively. The cosine similarity takes values between 0 and 1. This quantity is 1 if *i* had contacts in both days with exactly the same individuals, spending the same fraction of time in proximity of each one, and 0 if, on the contrary, *i* had contact in day 2 with totally different persons with respect to day 1.

#### Analysis of contacts between infant and other household members

For each household, and for each household member, we calculated an Infant Proximity Score (IPS) describing the contact pattern between the infant and other family members. The IPS was defined as the fraction of time spent by each family member in proximity with the infant, over the total time spent by the infant at home.

In other words, the IPS quantifies the fraction of time spent by the infant at home and in contact with another household member, therefore, it represents the proportion of time spent at home, in which the child is at a potentially higher risk of infection transmission by a potentially infected household member. The normalization over the time spent at home by each infant, allows comparability among different families.

The Ordinary Least Squares (OLS) regression was used to investigate the effects of sociodemographic variables and of infant feeding habits on the observed contact patterns. In order to allow comparison of contact patterns between families, we used only the maternal and paternal IPS in the regression model, since other family members were not present in all the households. The standard regression method based on the OLS estimates the effect of explanatory variables at the conditional mean of the distribution of the dependent variable. Socio-demographic variables and feeding habits were included in both models as explanatory variables. The explanatory variables were categorised as follows: parents’ age ≤ 35 or > 35 years; parents’ sex; infant’s age 0-3 or 4-6 months, presence of siblings: yes/no; infant’s feeding: exclusive breastfeeding vs partial breastfeeding/artificial feeding. Statistical analyses were carried out using the R software, version 3.1.2 [25]; statistical significance was set at a p < 0.05 level.

## Results

### General description of the study population

We enrolled a total of 20 households, and a code was assigned to each family. A total of 4 families and one member of one family were excluded from the analysis. Three of the first enrolled families (H04, H05 and H08) were excluded from the analysis due to an human error in the sensors’ deployment. Specifically, in the first phase of the enrolment process, the devices were all activated and included in the plastic pouches at the same time and were only later distributed to the families, leading to battery discharge before the data collection. To avoid such issue, in all subsequent deployments the devices were activated only right before being given to the enrolled families, allowing for a longer battery duration.

Household H19 was excluded because the contacts measured by sensors did not correspond to the times of entry and exit from the house indicated by family members.

Finally, data collected by the sensor assigned to the father of H02 were not included in the analysis because the recorded contacts did not correspond to the reported times of entry and exit from the house. Therefore, in H02, we considered only the contact data between the mother and the infant.

Overall, 16 households were included in the data analysis, accounting for a total of 55 sensors, distributed as follows: 16 infants, 4 pre-school children, 31 adults (16 mothers, 15 fathers) and 4 elderlies (1 grandfather, 3 grandmothers). The average household size was 3.44. One family had 2 family members (infant and mother), 9 families had 3 family members (infant, mother and father), 5 families had 4 family members (in 2 families: infant, mother, father, and grandparent; in 3 families: infant, mother, father, and sibling), 1 family had 6 family members (infant, mother, father, sibling, and 2 grandparents). In 4 families (25%), the infant had siblings, and in 3 families (18.75%) grandparents lived in the same house. The median infant’s age was 3.4 months (average = 3.6; range 1.8 - 5.9), 10 (62.5%) of infants received exclusive breastfeeding, 5 (31.25%) received partial breastfeeding and 1 (6.25%) received artificial feeding. None of the infants attended a nursery or other communities. The infants spent at home an average of 19.66 hours (median = 19.18; range 14.8 - 23.5), corresponding to 81.94% of the 24 hours. In our sample, all infants spent the night at home for the whole study period.

### Contact data and network analysis among all participants

Figure 1 shows the probability distribution of person-to-person contact durations for all individuals over the whole experimental period. The average contact duration measured on all contact events was 200.12 seconds. We recorded 5,958 contact events over the whole experimental time period. A total of 4,201 (70.51%) contacts were recorded between the infant and other family members, with a daily average of 79 contacts (range 6 - 133). Mean duration of contacts between infants and other family members was 234.92 seconds. We generated contact matrices based on duration of contacts (in seconds) and on number of contacts (contact events) by age category (Figure 2) and by role (Figure 3). We considered as roles: infant, mother, father, sibling, and grandparent. We divided the total daily contact durations and the total daily number of contacts by the time during which two tags were simultaneously in the house, and by the total number of persons belonging to an age category and to a role, thereby obtaining the daily average of the contact durations and the daily average of contact events per capita. The highest average contact durations and average number of contacts corresponded to the contacts between infants (<6 months old) and adults (20-49 years old, *i.e.,* parents), in particular between infant and mother.

**Figure 1.**
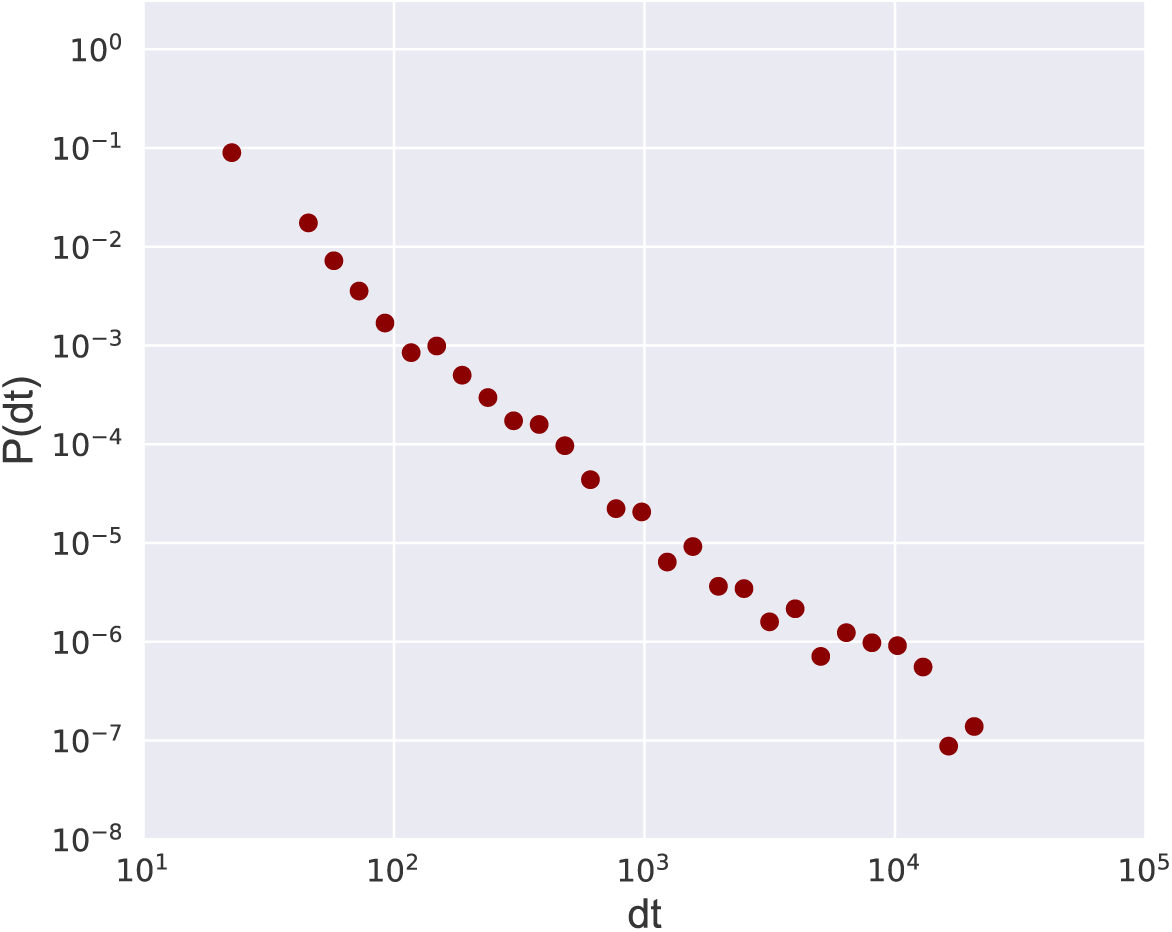
Distribution of contact duration. Probability density distribution P(dt) of contact duration dt measured for all individuals over the whole experimental period.

**Figure 2.**
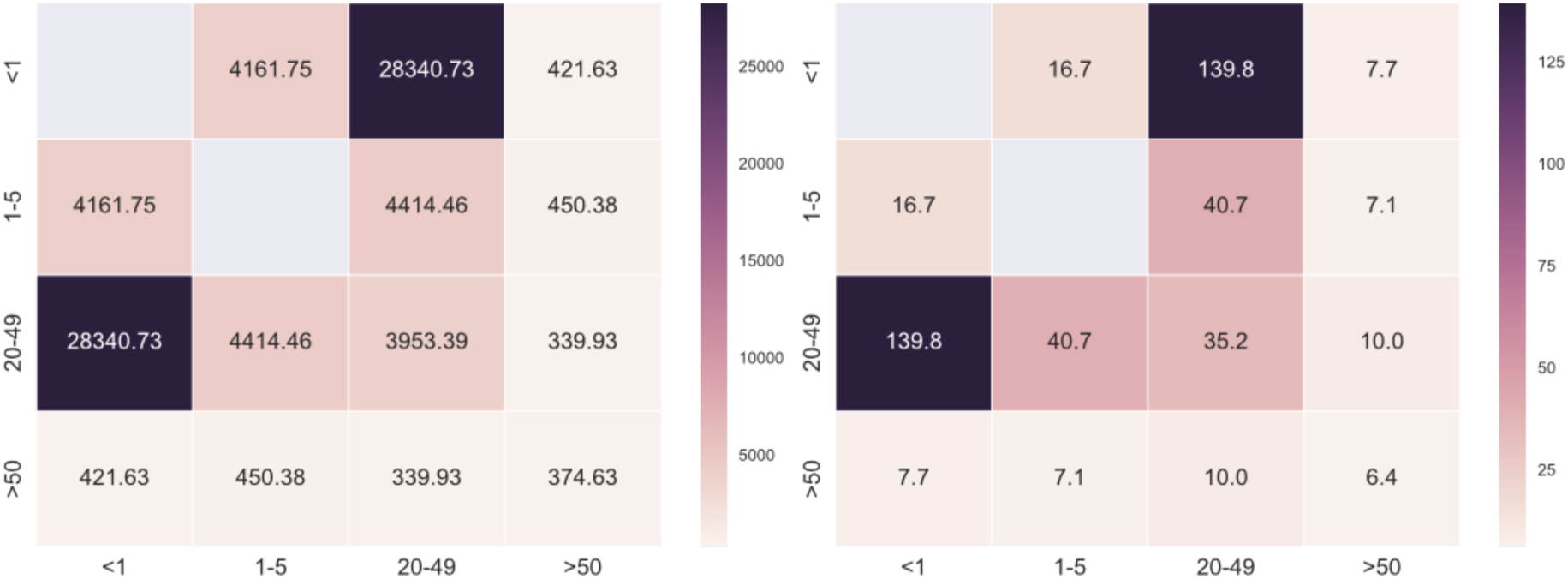
Contact matrices of the average contact durations *per capita* (left panel) and the average contact events *per capita* (right panel) by age categories for all individuals.

**Figure 3.**
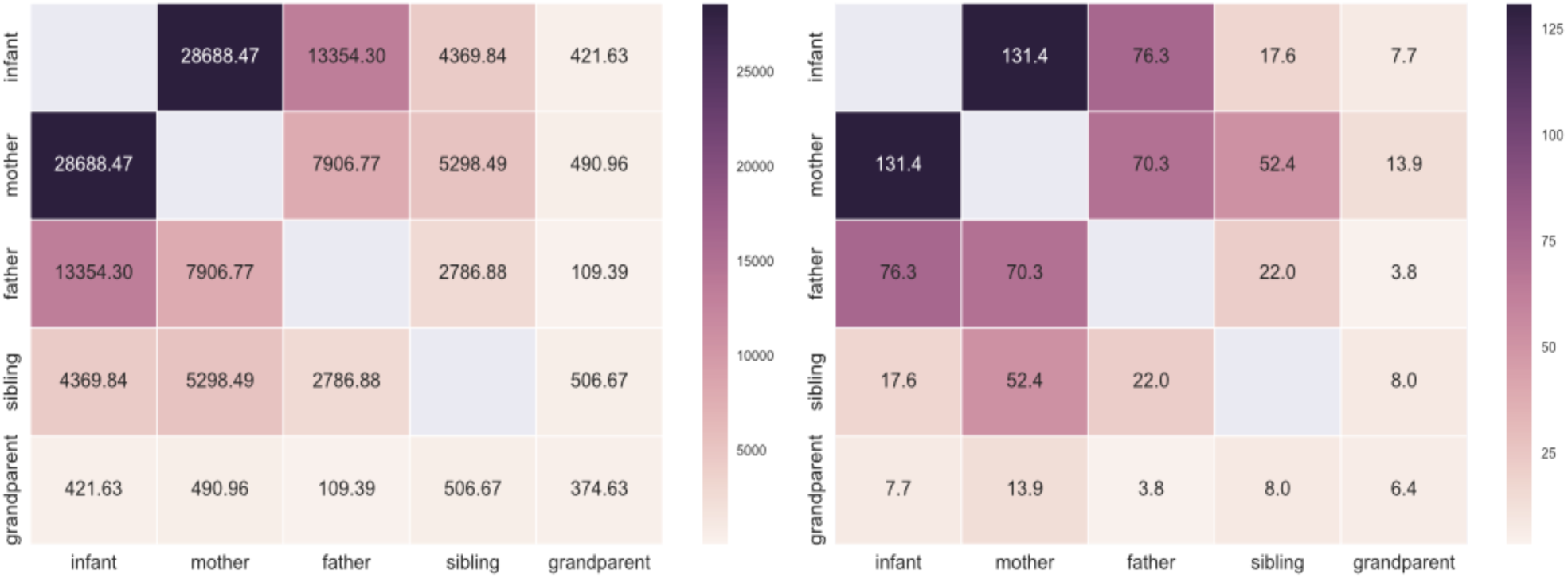
Contact matrices of the average contact durations *per capita* (left panel) and the average contact events *per capita* (right panel) by roles for all individuals.

Figure 4 displays a representation of the contact network of the 16 households, where nodes are individuals and edges indicate the presence of at least one recorded contact during the study period. Nodes are color coded according the role of the family members, and the edge thickness is proportional to the weight w_ij_, that is the total amount of time spent in proximity by two individuals. Edge’s color indicates the time of the day during which it was mainly active. Overall, 45.7% of the total time spent in proximity by participants is observed during the morning, 14.4% in the late morning – lunch time, 20.2% in the afternoon, and 19.7% in the late afternoon - evening. The overall contact network is formed by 55 nodes and 69 edges, each single connected component representing a household. The network density is 1.0 for all households except for H07, H09, H12, and H20, where the density ranged between 0.67 and 0.83.

**Figure 4.**
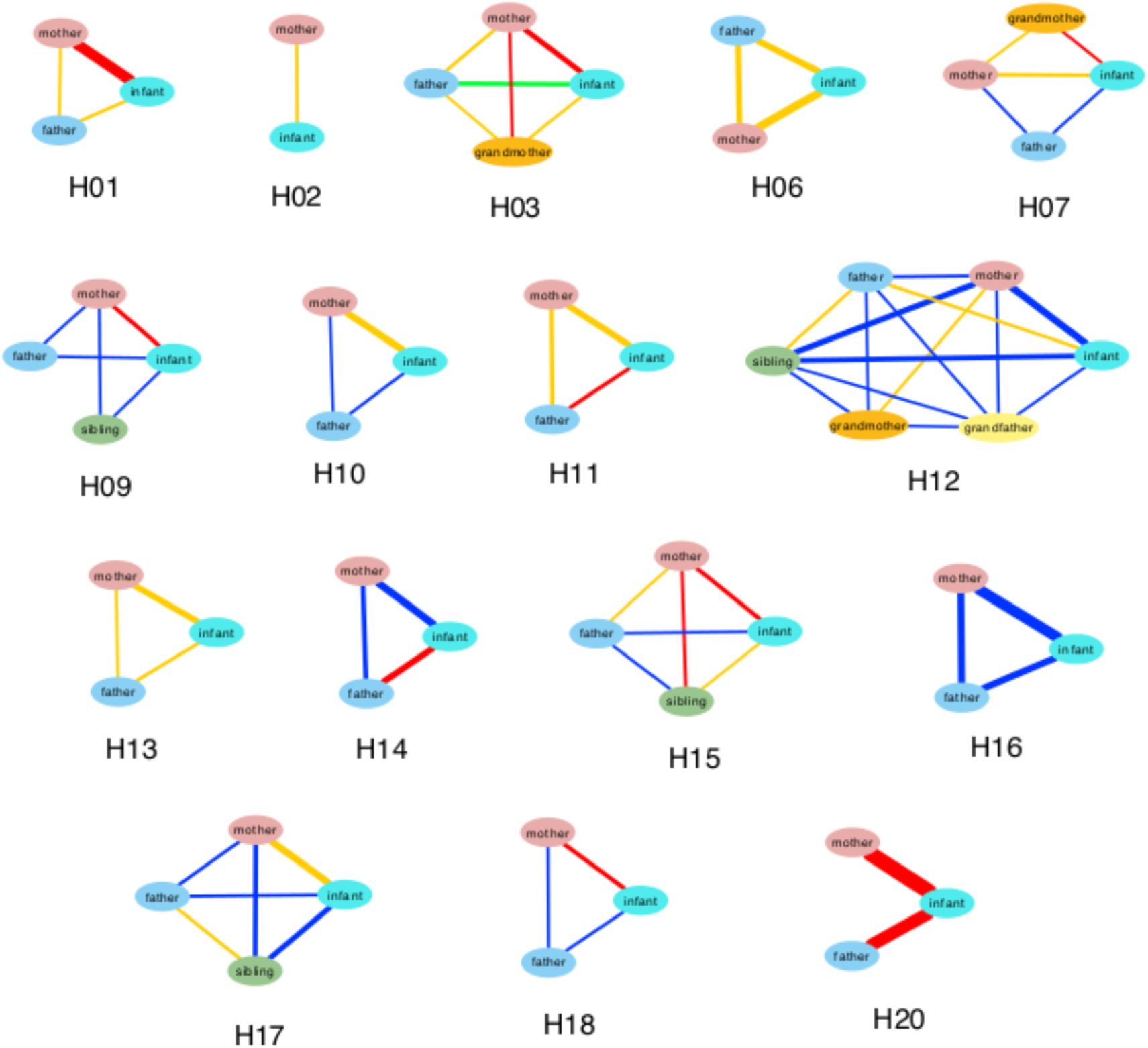
Contact networks between household members aggregated over the whole study duration. Nodes are color-coded according to the role of the family member. Edge thickness is proportional to the total time spent in proximity by two connected individuals, and the edge color indicates the time of the day during which it was mainly active: *blue* = morning, *yellow* = late morning - lunch, *red* = afternoon, and *green* = late afternoon - evening.

Daily aggregated contact networks were generated by aggregating all the contact events measured from 7 am to 9 pm for each household. To evaluate the changes in the daily contact durations of each household member, we computed the cosine similarity between the neighbourhoods of each node in each pair of daily networks. The distributions of similarities are shown in Figure 5, aggregated for all households and by each household. The households H02, H10, H15 and H17 were not included in this analysis because their contacts were not measured for more than one entire day (from 7 am to 9 pm), and data were not comparable. The median values of the cosine similarity all varied between 0.7 and 1, while the average network density varied between 0.67 and 1, indicating a substantial stability of individual contact patterns across days.

**Figure 5.**
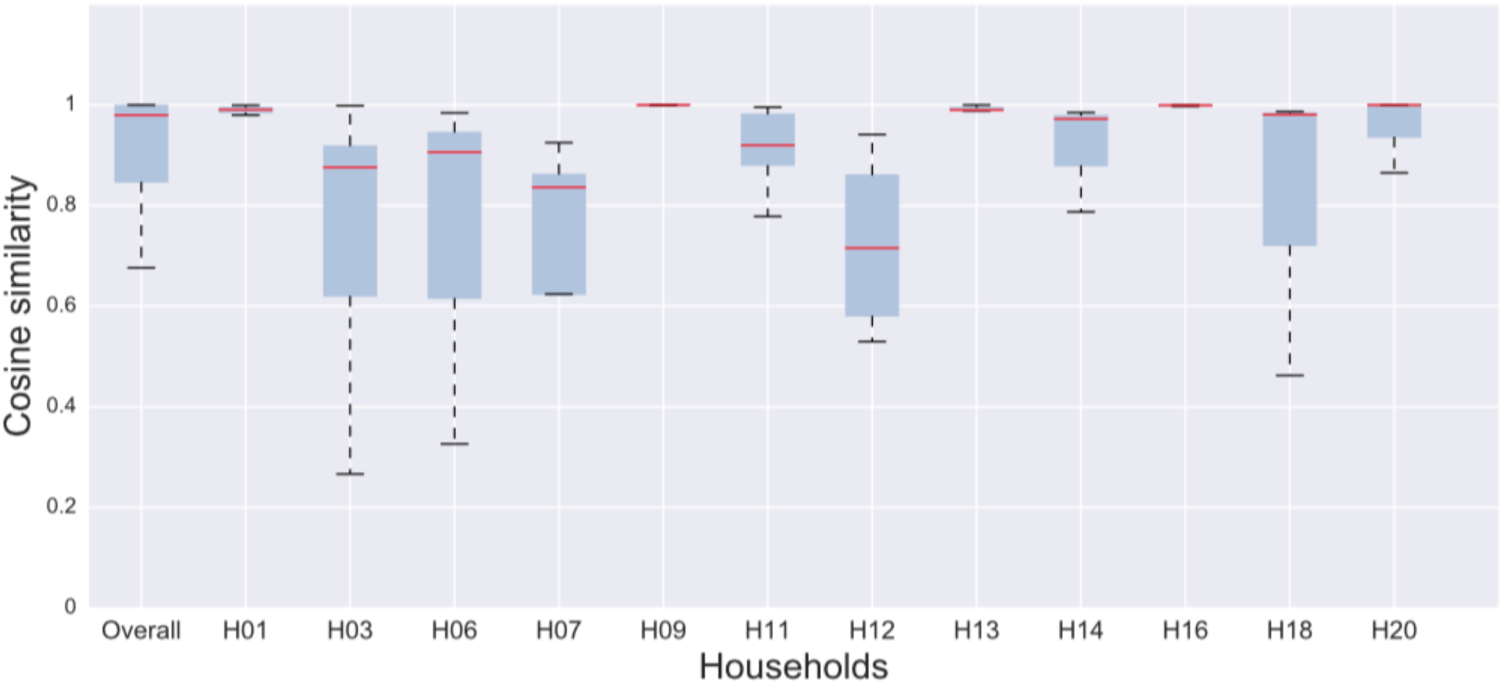
Distribution of cosine similarity measured for each node of the full contact network (overall) and for each single household. Distributions are obtained by measuring the cosine similarity of each node's neighbourhood, for each pair of days of data collection. The box and whisker plots show the interquartile range, and the red line indicates the median value. The error bars extend from the box to the highest and lowest values.

### Contact patterns between infant and other household members

For each household, we calculated the infant proximity score (IPS) between the infant and each family member, defined as the percentage of time spent by each family member in proximity to the infant, over the total time spent by the infant at home (Figure 6). The average IPS was 18.55 (median = 14.21; range 0.51 - 77.52) considering all family members.

**Figure 6.**
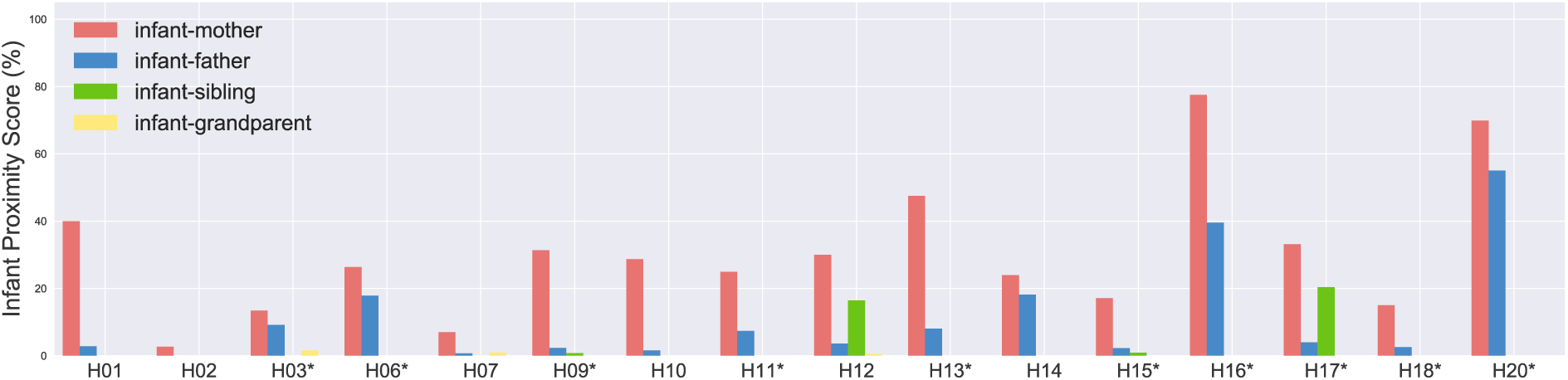
Infant Proximity Score (IPS) between each family members and the infant (IPS = percentage of time spent by each family member in proximity to the infant, over the total time spent by the infant at home). *Pink* = IPS between mother and infant; *blue* = IPS between father and infant, *green* = IPS between sibling and infant; *yellow* = IPS between grandparent and infant. Families in which the infant is exclusively breastfed are labelled with *.

Then, we used an Ordinary Least Squares (OLS) regression to investigate the effects of sociodemographic variables and feeding habits on the parents’ IPS (average = 21.40; median = 17.09; range 0.72 - 77.52). The results showed that the IPS was most affected by infant’s feeding habit, parent’s sex and infant’s age (F = 5.057; R-squared = 0.403; p = 0.002). The IPS was significantly higher for exclusive breastfeeding (p = 0.043), and for infants aged 4-6 months (p = 0.038), and it was significantly lower for fathers (p = 0.002) (Table 1).

**Table 1.** Results of the OLS regression predicting the effects of infant’s feeding (exclusive breastfeeding vs partial breastfeeding/artificial feeding), parents’ sex, presence of siblings (yes/no), parents’ age (≤ 35 or > 35 years), and infant’s age (0-3 or 4-6 months) on parents’ Infant Proximity Score (IPS). The value of the estimate refers to the variable in brackets.

| **Variable** | **Estimate** | **Standard Error** | ***t*-value** | ***p*-value** |
| --- | --- | --- | --- | --- |
| *Intercept* | 15.132 | 7.384 | 2.049 | 0.051 |
| *Infant's feeding (exclusive breastfeeding)* | 12.808 | 6.003 | 2.134 | 0.043 |
| *Parents' sex (M)* | -20.440 | 5.836 | -3.502 | 0.002 |
| *Presence of siblings (yes)* | -9.565 | 6.403 | -1.494 | 0.148 |
| *Parents' age (> 35 years)* | 7.035 | 6.048 | 1.163 | 0.256 |
| *Infant's age (4-6 months)* | 12.991 | 5.946 | 2.185 | 0.038 |

## Discussion

With the present study, we report the first quantitative assessment of social contact patterns in households with infants younger than 6 months, based on a technology that allows for high-resolution measurements of social contacts. We obtained contact matrices that can have a crucial role for understanding transmission dynamics and for informing strategies aimed at the prevention of infectious diseases in very young infants. Our study shows that using wearable sensors based on RFID technology is feasible for obtaining a precise measurement of the pattern of close contacts among individuals living in the same house. We quantitatively demonstrated that 1) compared to social contacts of children in the age group 0-5, the contact patterns of an infant <6 months within a household is unique and not assortative, in agreement with previous results of a diary-based study [26]; 2) the mother has significantly more frequent and longer contacts with the infant, compared to the father and to other family members, thus being the main potential source of infection; nevertheless, the contribution of other family members to the infection risk is not negligible. Finally, our results suggest exclusive breastfeeding may be associated to higher contact frequency and duration.

Infants spent almost all of their time (80%) inside the house, together with other household members. A strong cultural and seasonal variability may exist for this kind of figure. Nevertheless, taking into account that respiratory infections spread more easily in close environments, our results suggest that prevention strategies dedicated to infants should primarily focus on the transmission risk in this kind of context.

We assessed the similarities between contacts measured among household members on different days, keeping in mind the constraints imposed by the short study duration. The results obtained by the cosine similarity showed that the durations of time spent in proximity with different family members were similar from day to day, in agreement with what observed by Kiti *et al.* [24] in a similar sensor study set in a rural African context, and with a recent contact survey study in Belgian households [27]. These findings suggest that a few days of data collection could be adequate for an accurate description of contact patterns within households, as contacts are stable and repetitive across single days.

The analysis of temporal features of contact links between household members revealed a variability in the time of the day during which two individuals mainly interacted, although most contacts took place during morning and at lunchtime, when household members congregate for breakfast and lunch [24].

Previous survey studies have measured the contact patterns of infants both within and outside households. Van Hoek *et al.* [26] reported an average daily contact frequency of 6 for infants younger than 10 weeks and of 7.6 for infants 10 weeks-12 months of age. Mossong *et al.* [28], in the original article reporting results from the POLYMOD study, reported an average 10.2 contacts for the 0-4 years age group (contact patterns within this age group could actually be very heterogeneous). A recent study by Campbell *et al.* [29] estimated the contacts of infants younger than 1 year in three Australian communities by surveying the contact patterns of their primary carers (the mother) and found a mean daily number of contacts of infants ranging between 6.5 and 9.1.

In our study, the average degree of the infants in the aggregated contact network was equal to 2.37 (i.e. 2.37 unique contacts over the study duration), thus in line with the above findings, considering that we did not measure contacts with non-household members and that the average household size was 3.4. However, when comparing the results of sensor-based studies, like ours, to paper-based contact diaries, it is important to consider the differences between the two methodologies which can be ascribed to different underlying contact definitions, under- and over-recording with sensors, and under- and over-reporting with contact diaries [30]. For instance, both the study by Van Hoek *et al.* [26] and the study by Mossong *et al.* [28], as well as other diary-based studies [31], adopted a specific definition of contact event, *i.e.* an either a skin to skin contact or an interaction in close proximity with three or more words (directed to the infant or in a two-way conversation). In our study, a contact was recorded when an interaction < 1.5 m occurred, likely yielding a higher frequency of contacts compared to the conversation/skin-to-skin definition. More in general, survey based studies measure the number of unique contacts, thus missing the number of repeated interactions between study participants, which are instead captured by proximity sensors. Secondarily, the diary-based studies might have underestimated the true contact frequency due to recall bias. In diary based studies, the data collection method (*e.g*., paper vs online diary) has been shown to influence the reported contact frequency [31], suggesting that recall bias may play an important role in the quantification of daily contact events. While the conversation/skin-to-skin definition might be more specific regarding the infection transmission risk, the RFID measurement allows to objectively quantify the contact events, and the 1.5 m threshold is sensitively representative of the risk of transmission by droplet for respiratory infections [32].

The contact networks showed a high heterogeneity of the cumulative time spent in proximity with the infant by family members, despite the small number of individuals and the short duration of the study. Most of the contacts occurred between the infant and other family members (70%), and, as showed by the contact matrices, many contacts were observed between infants and adults, in particular between infant and mother, followed by father, siblings and grandparents. Our results show a highly heterogeneous distribution of contact durations characterized by a heavy tail: most contacts are short and there are few long-lasting contacts, whose probability is, however, not negligible. A similar broad distribution of contact durations has been observed in other settings, including schools [11], hospitals [8], and rural households [24].

Siblings and grandparents were scarcely represented in our sample of households; their contribution to the infection risk might be limited, compared to the probability of transmission from parents. We enrolled only individuals living in the same house of the infant. Thus, we might have underestimated the contribution to the contact pattern by grandparents visiting the infant from outside the house. Despite such potential bias, grandparents living in the same household of the infant had 7.7 average contacts per day with the infant, for an average total duration of 421 seconds. Siblings were found to be in close contact with the infant more often compared to grandparents (17.6 contacts per day), with a much higher average contact duration (4,369 s). These figures, anyway, are distinctly lower when compared to contact patterns between the parents and the infant (76 contacts per day for fathers, with a 13,354 s average contact duration, and 131 contacts per day for mothers, with a 28,688 s average contact duration).

Our results may have implications for vaccination strategies. First of all, we showed that the contacts between the infant and the mother have a remarkably higher frequency and duration compared to contacts between the infant and other household members. The mother has been demonstrated to be the main source of pertussis infection for the infant [15, 33, 34] and for other family members [16]. Moreover, the cocooning strategy aimed at protecting young infants from pertussis through the vaccination of household members may be ineffective and difficult to implement [35]. Taking into account these observations, our results represent a further evidence in favour of maternal vaccination against pertussis during pregnancy. This strategy, targeting the crucial nodes of the contact networks in infant households (*i.e.* the mother and the infant) has been repeatedly demonstrated to be safe [36], effective [37] and cost-effective [38].

Families in which the baby is exclusively breastfed tend to have a more intense contact pattern compared to families in which the infant receives artificial of mixed feeding. The implication of this observation is delicate and controversial. Breastfeeding is universally recognised to have a protective effect towards infections, by means of a variety of immune properties. The WHO recommends to always continue breastfeeding if the mother has an infection, with very few exceptions (*e.g.,* HIV and HTLV infections). Most studies show that exclusive breastfeeding reduces the risk of infection if it is continued for longer than 4 or 6 months [39], and, often, the assessment of the risk reduction is performed on children older than 1 year; data on the infection risk in the first months of life is lacking. In our study, we show that breastfeeding is associated with a more intense contact with parents. This could actually imply a higher risk of infection transmission. Although the infection risk is probably balanced by the anti-infective properties of breast milk, our results suggest that a cautious approach should be used if the mother or any other family member is sick, as clearly pointed out in the CDC Guidance for the Prevention and Control of Influenza in the Peri- and Postpartum Settings [40]: the sick family member should frequently wash her or his hands and wear a mask when in close contact with the infant.

It is important to highlight some key limitations of the present study. Contact patterns are influenced by cultural and demographic differences across countries [41, 42]. Our assessment was focused on a limited group of families living in a specific area of Rome, Italy, attending the office of a single family paediatrician. To obtain more reliable contact matrices for informing transmission models, a larger and more representative national/international sample would be desirable, and the study design should take into account a longer period of data collection for each family. On the other hand, the methods presented in this paper can be useful to detect contact patterns in very specific contexts, thus allowing implementing tailored prevention strategies accordingly.

We considered in the analysis only data collected during daytime and inside the house. All night contacts (collected from 9 pm to 7 am), were disregarded during the analysis, nevertheless, families have different sleeping habits and infants often awake at night, therefore, monitoring social interactions during the night would have probably provided additional information regarding the infection risk. At the beginning of the enrolment, an issue regarding the device preparation compromised some of the tags’ functions. While it may not be possible to detect tag malfunctions during the data collection, future studies could minimise human error by a proper training on how to prepare and use the sensors, especially when collecting data for longer periods.

In conclusion, with the present study, we demonstrated the feasibility of measuring mixing matrices within infants’ households using wearable proximity sensing devices. A system based on RFID technology may overcome the recall bias which is intrinsic to diary-based studies on contact patterns, in particular in the case of young infants, whose behaviour can only be indirectly monitored by a guardian or someone who spends most of the time with the infant [43].

Moreover, our results suggest that, as the mother is responsible for the large majority of the infant’s contact pattern, and as the contribution to the contact pattern by other family members is very variable, vaccination against pertussis during pregnancy is probably the best strategy to protect young, unvaccinated infants [37]. A further study, involving a larger and more representative sample of families would be advisable to obtain more robust contact data.

## Acknowledgements

This study was supported by the grant n. RF-2010-2317709 from the Italian Ministry of Health. This study was supported by the Lagrange Project of the ISI Foundation funded by the CRT Foundation to LO, MT and CC.

## Supporting Information

**Table S1.** Number of family members, sex and age distribution, time start and time stop of the experiment.

|  |  | Sex |  | Age categories |  |  |  | Experimental period |  |
| --- | --- | --- | --- | --- | --- | --- | --- | --- | --- |
| Households | Number of family members | F | M | <6 months | 1-5 years | 20-49 years | Age >=50 | Time start | Time stop |
| *Included in data analysis* |  |  |  |  |  |  |  |  |  |
| H01 | 3 | 2 | 1 | 1 | 0 | 2 | 0 | 03/03/15<br>07:00 | 06/03/15<br>22:00 |
| H02 | 2 | 1 | 1 | 1 | 0 | 1 | 0 | 24/03/15<br>07:00 | 28/03/15<br>22:00 |
| H03 | 4 | 2 | 2 | 1 | 0 | 2 | 1 | 14/04/15<br>13:45 | 18/04/15<br>20:00 |
| H06 | 3 | 1 | 2 | 1 | 0 | 2 | 0 | 29/10/15<br>13:40 | 02/11/15<br>00:00 |
| H07 | 4 | 3 | 1 | 1 | 0 | 2 | 1 | 03/11/15<br>11:00 | 06/11/15<br>12:00 |
| H09 | 4 | 2 | 2 | 1 | 1 | 2 | 0 | 11/11/15<br>16:20 | 14/11/15<br>12:00 |
| H10 | 3 | 2 | 1 | 1 | 0 | 2 | 0 | 01/12/15<br>12:00 | 04/12/15<br>00:00 |
| H11 | 3 | 1 | 2 | 1 | 0 | 2 | 0 | 01/12/15<br>12:30 | 05/12/15<br>23:00 |
| H12 | 6 | 4 | 2 | 1 | 2 | 2 | 2 | 01/12/15<br>13:00 | 04/12/15<br>14:00 |
| H13 | 3 | 2 | 1 | 1 | 0 | 2 | 0 | 02/12/15<br>17:50 | 05/12/15<br>16:15 |
| H14 | 3 | 2 | 1 | 1 | 0 | 2 | 0 | 17/12/15<br>14:00 | 20/12/15<br>23:00 |
| H15 | 4 | 1 | 3 | 1 | 1 | 2 | 0 | 17/12/15<br>15:30 | 20/12/15<br>23:30 |
| H16 | 3 | 1 | 2 | 1 | 0 | 2 | 0 | 18/12/15<br>08:45 | 21/12/15<br>12:30 |
| H17 | 4 | 2 | 2 | 1 | 1 | 2 | 0 | 21/12/15<br>18:00 | 24/12/15<br>19:00 |
| H18 | 3 | 1 | 2 | 1 | 0 | 2 | 0 | 28/12/15<br>19:35 | 31/12/15<br>20:10 |
| H20 | 3 | 1 | 2 | 1 | 0 | 2 | 0 | 08/01/16<br>15:00 | 13/01/16<br>20:00 |

| *Excluded from data analysis* |  |  |  |  |  |  |  |  |  |
| --- | --- | --- | --- | --- | --- | --- | --- | --- | --- |
| H02 | 1 | 0 | 1 | 0 | 0 | 1 | 0 | 24/03/15<br>07:00 | 28/03/15<br>22:00 |
| H04 | 4 | 1 | 3 | 1 | 1 | 2 | 0 | 16/04/15<br>18:30 | 19/04/15<br>20:00 |
| H05 | 3 | 2 | 1 | 1 | 0 | 2 | 0 | 17/04/15<br>09:00 | 21/04/15<br>22:00 |
| H08 | 3 | 1 | 2 | 1 | 0 | 1 | 1 | 03/11/15<br>15:30 | 04/11/15<br>18:00 |
| H19 | 5 | 2 | 3 | 1 | 2 | 2 | 0 | 05/01/16<br>10:00 | 07/01/16<br>21:00 |

**Table S2.** Network statistics of each household: number of nodes, number of edges, average degree, network density, and average daily network density.

| Households | Number of nodes | Number of edges | Average degree | Network density | Average daily density |
| --- | --- | --- | --- | --- | --- |
| H01 | 3 | 3 | 2.0 | 1.0 | 1.0 |
| H02 | 2 | 1 | 1.0 | 1.0 | - |
| H03 | 4 | 6 | 3.0 | 1.0 | 1.0 |
| H06 | 3 | 3 | 2.0 | 1.0 | 1.0 |
| H07 | 4 | 5 | 2.5 | 0.83 | 0.83 |
| H09 | 4 | 5 | 2.5 | 0.83 | 0.83 |
| H10 | 3 | 3 | 2.0 | 1.0 | - |
| H11 | 3 | 3 | 2.0 | 1.0 | 1.0 |
| H12 | 6 | 14 | 4.7 | 0.93 | 0.77 |
| H13 | 3 | 3 | 2.0 | 1.0 | 1.0 |
| H14 | 3 | 3 | 2.0 | 1.0 | 1.0 |
| H15 | 4 | 6 | 3.0 | 1.0 | - |
| H16 | 3 | 3 | 2.0 | 1.0 | 1.0 |
| H17 | 4 | 6 | 3.0 | 1.0 | - |
| H18 | 3 | 3 | 2.0 | 1.0 | 1.0 |
| H20 | 3 | 2 | 1.33 | 0.67 | 0.67 |

### Degree and weight distributions

The average degree of the contact network is 〈k〉 = 2.5, and the degree distribution extends between kmin = 1 and kmax = 5. Figure S1 shows the cumulative degree distribution of the contact network aggregated for all individuals over the whole experimental period (left panel) (*i.e.,* probability that a randomly chosen node has degree k), and the distribution of the weights of the aggregated contact network (right panel).

**Figure S1.**
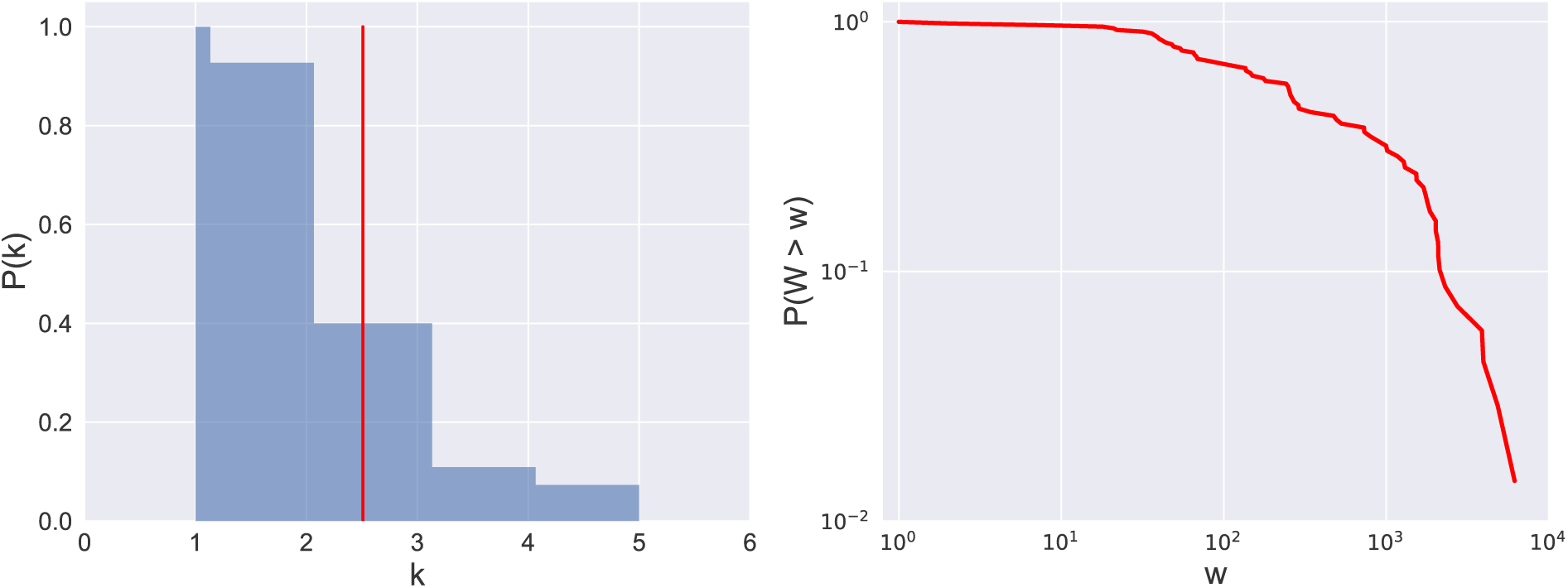
Statistics of the aggregated contact network. Cumulative degree distribution P(k) of the contact network aggregated for all individuals over the whole experimental period. The red line indicates the average degree value, 〈k〉 = 2.5. (left panel). Complementary Cumulative Distribution Function (CCDF) of weights (i.e. cumulated contact durations) of the aggregated contact network (right panel).

